# A Context-Aware Single-Cell Proteomics Analysis pipeline

**DOI:** 10.64898/2026.04.03.716382

**Authors:** Carla Salomó Coll, Agata N Makar, Alejandro J Brenes, Joseph Inns, Matthias Trost, Neil Rajan, Simon Wilkinson, Alex von Kriegsheim

## Abstract

Single-cell proteomics (SCP) by mass spectrometry can now quantify hundreds to thousands of proteins per cell, but the field still lacks standardised analytical pipelines that accommodate the diversity of instruments, sample preparation workflows and biological contexts encountered in practice. Existing workflows, largely adapted from single-cell transcriptomics, do not account for the informative missingness, pervasive ambient protein contamination and limited feature space that distinguish proteomic from transcriptomic data. In addition, cell type annotation remains a manual bottleneck that is subjective, difficult to reproduce and hard to scale.

Here we present an end-to-end pipeline that integrates adaptive quality control, entropy-guided iterative batch correction, multi-modal marker discovery that exploits detection patterns unique to proteomics, and context-aware annotation by large language models (LLMs) coupled to structured contradiction reasoning and orthogonal data-driven validation. Benchmarking on published single-cell proteomic datasets from developing human brain and glioblastoma-associated neutrophils revealed systematic LLM failure modes, including context-insensitive marker vocabulary and misinterpretation of phagocytic or lytic cell states. We addressed these errors using a three-round prompt architecture that combines general biological principles with auto-generated dataset-specific constraints.

In held-out validation on a skin tumour dataset acquired, the pipeline showed high concordance with FACS-sorted ground truth. In the caerulein-injured pancreas, orthogonal immunohistochemistry further supported annotations of macrophage, stellate and immune populations. The pipeline is fully automated under fixed settings, and available as Context-Aware Single-Cell Proteomics Analysis (CASPA), providing SCP laboratories and facilities with a reproducible workflow that delivers interpretable, confidence-quantified annotations suitable for downstream expert review.

## Introduction

Single-cell proteomics (SCP) enables direct measurement of protein abundance at single-cell resolution, avoiding the well-recognised discordance between mRNA and protein levels that can limit transcriptomic inference in many biological settings ^1, 2^. Advances in ultra-low-input mass spectrometry now allow quantification of hundreds to thousands of proteins from individual cells ^3, 4^. However, while the underlying measurement technologies have progressed rapidly, the analytical infrastructure required to interpret these data has lagged behind. As a result, an increasing proportion of SCP datasets are generated by technology-focused laboratories and associated core facilities that may not have the specialist biological or bioinformatic expertise needed to deliver interpretable results to downstream collaborators.

Most current SCP analysis workflows are adapted from single-cell RNA-seq toolkits developed for high-dimensional transcriptomic matrices ^5^. This is problematic because SCP data differ from transcriptomic data in several fundamental ways. First, sparsity is informative: failure to detect a protein may reflect genuine biological absence, technical dropout or ambient carryover, and these scenarios have distinct analytical implications ^6^. Second, batch effects are common in multi-batch SCP experiments, yet most workflows treat batch correction as a one-step procedure without explicitly evaluating whether correction is adequate ^7^. Third, marker discovery is often based on a single statistical framework, even though detection patterns, intensity differences, model-corrected effects and pathway-level signatures each capture complementary aspects of cell identity in proteomic data. Together, these limitations mean that SCP analysis still depends heavily on manual intervention, reducing both throughput and reproducibility.

Cell type annotation remains a particular bottleneck. Reference-based classifiers trained on transcriptomic atlases generally perform poorly on proteomic data, where canonical markers are often not detected, and the feature space is substantially smaller ^8^. Manual annotation by domain experts remains flexible and effective, but it is inherently subjective and difficult to scale. Large language models (LLMs) have recently shown promise for biological interpretation ^9^. However, when applied naively, they can generate non-deterministic outputs and hallucinated reasoning.

Here, we present a robust end-to-end analysis pipeline for SCP designed to address these limitations. The workflow combines adaptive quality control, iterative batch correction, multi-modal marker discovery and a three-tier annotation framework that integrates context-aware LLM reasoning with orthogonal data-driven validation. We assess the pipeline across four datasets of increasing analytical difficulty and tissue origin; The developing human brain, glioblastoma-associated neutrophils, skin tumours and caerulein-injured pancreas. Across these settings, we observe strong concordance with published expert annotations, identify systematic LLM failure modes and define practical limits for automated annotation in single-cell proteomics.

## Results

### An end-to-end analytical framework for single-cell proteomics

We developed an automated analysis pipeline that converts raw protein-group matrices into quality-controlled, batch-corrected, clustered and annotated single-cell proteomics datasets (Fig. 1a). The workflow accepts output from the major DIA search engines, DIA-NN and Spectronaut^10, 11^, and was designed around three practical requirements: adaptation to heterogeneous datasets, integration of complementary analytical modalities and tiered validation. This reflects the reality that SCP data vary substantially across instruments, sample preparation workflows and biological contexts, and that no single analytical modality is sufficient for robust interpretation. A complete workflow schematic, parameter summary and statistical overview are provided in Supplementary Fig. 1.

**Figure 1.**
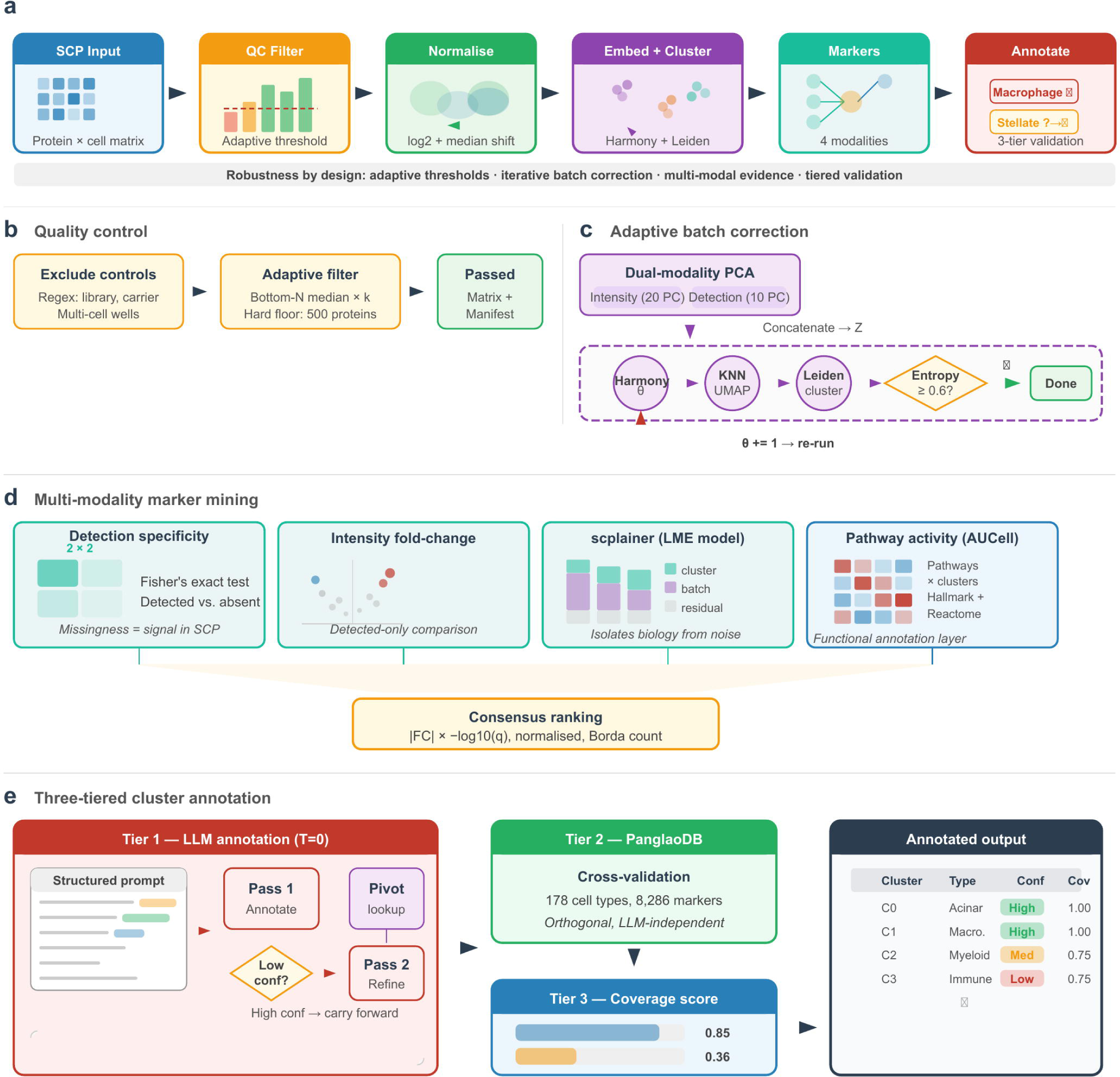
Robust end-to-end pipeline for single-cell proteomics analysis. (**a**) Pipeline overview. Raw protein group matrices from any standard search engine are processed through six stages: quality control with adaptive cell filtering, log2 transformation and per-cell median normalisation, joint embedding with iterative batch correction, multi-modality marker discovery, and three-tiered cell type annotation. (**b**) Quality control module. Cells are filtered using an adaptive threshold that adjusts to dataset quality distribution, with a configurable hard floor. A novel cluster-per-batch diagnostic flags batches with abnormal cluster composition. (**c**) Adaptive Harmony batch correction. A dual-modality PCA jointly embeds protein intensity and detection patterns. The Harmony diversity penalty (θ) is iteratively increased until batch mixing entropy reaches the target threshold (default: 0.6) or the maximum θ is reached, ensuring adequate correction without manual parameter tuning. (**d**) Multi-modality marker mining. Four complementary approaches - detection specificity (Fisher’s exact test), intensity fold-change (Mann-Whitney U, detected-only), scplainer linear model (variance decomposition into cluster, batch, and residual components), and pathway activity (AUCell) - are integrated through normalised Borda consensus ranking. (**e**) Three-tiered cluster annotation. Tier 1: a context-aware LLM annotates clusters in two passes, with automated supplemental marker queries for ambiguous clusters. Tier 2: PanglaoDB cross-validation provides orthogonal, LLM-independent assessment. Tier 3: quantitative marker coverage score anchors confidence in the observed data. The final output includes cell type labels, confidence tiers, supporting and contradicting markers, resolving markers for follow-up experiments, and a full audit trail.

The pipeline begins with adaptive quality control rather than fixed filtering rules (Fig. 1b). Control runs are removed automatically, and remaining cells are filtered using a dataset-specific threshold derived from the lower tail of the observed protein-count distribution or annotated blank/empty wells. In addition, we employ a minimum threshold of 400 detected proteins as implemented previously ^12^. This avoids a common failure mode of fixed cut-offs, which can be too permissive for high-quality datasets and too stringent for shallower ones. In addition to standard per-cell metrics, the workflow reports a cluster-per-batch composition diagnostic that highlights batches whose cluster distributions diverge from the global pattern, enabling detection of technically compromised batches that would not necessarily fail cell-level quality control (Fig. 1b)

To address batch effects, the pipeline uses an iterative rather than single-pass correction strategy (Fig. 1c). Cell embeddings are constructed from a dual-modality principal component analysis (PCA) that combines protein intensity with binary detection patterns, leveraging the fact that in SCP, protein detectability can be as informative as abundance. After each Harmony iteration ^13^, batch mixing is quantified across Leiden clusters using weighted Shannon entropy on a 0-1 scale, and the diversity penalty is increased until the target mixing threshold is reached or a maximum setting is encountered. This design allows correction strength to adapt automatically to dataset complexity while preserving an explicit convergence criterion.

Cluster annotation integrates four marker modalities that capture complementary aspects of cell identity (Fig. 1d). Detection specificity is assessed by Fisher’s exact test on binary protein presence or absence. Intensity differences are evaluated by detected-only Mann-Whitney testing, avoiding zero-inflation bias from treating non-detected values as quantitative zeros. Model-based testing with scplainer ^14^ separates cluster-associated effects from technical factors, and AUCell pathway scoring provides a functional layer that is less sensitive to loss of individual canonical markers. These signals are summarised through a consensus ranking, such that proteins supported across multiple modalities are prioritised over modality-specific outliers.

Annotation then proceeds through three tiers of evidence (Fig. 1e). In the first tier, cluster-level evidence is assembled into a structured prompt containing experimental context, modality definitions, confidence rules and contamination-handling guidance, and is submitted to an LLM at temperature zero. Ambiguous clusters trigger an automated supplemental-marker query and are then re-annotated in a second pass. The second tier compares assigned labels with PanglaoDB ^15^ marker sets, and the third computes marker coverage against researcher-defined panels. Together, these components provide not only a label, but also a transparent record of supporting markers, contradictory evidence, confidence level and candidate markers for follow-up validation.

### Benchmarking across published datasets reveals strengths and limits of automated annotation

We first benchmarked the pipeline on the developing human brain SCP dataset reported by Wu et al. ^16^, which provides a demanding test of lineage-level annotation across prenatal cortical populations. We inferred batches from the date stamp in the sample name and identified 26 batches. We analysed the full protein-group matrix without pre-filtering erythroid or contaminant populations, retaining 819 cells after quality control. Because the dataset combines substantial technical heterogeneity, successful analysis depends critically on robust batch correction.

The adaptive Harmony loop required three iterations to achieve the target level of batch mixing, with mean Shannon entropy increasing from 0.53 to 0.62 as the diversity penalty increased from 2.0 to 4.0 (Fig. 2a,b). Before correction, the embedding was dominated by batch structure; after correction, biologically coherent clusters became apparent. Residual technical structure remained visible, however, in the cluster-per-batch composition diagnostic. In particular, batch 16, the largest batch in the experiment, was disproportionately represented in five clusters enriched for keratin and serum proteins but lacking coherent neural markers (Fig. 2c). Rather than discarding the entire batch, the pipeline localised likely technical compromise to these specific clusters, while preserving better-mixed neural populations for downstream interpretation.

**Figure 2.**
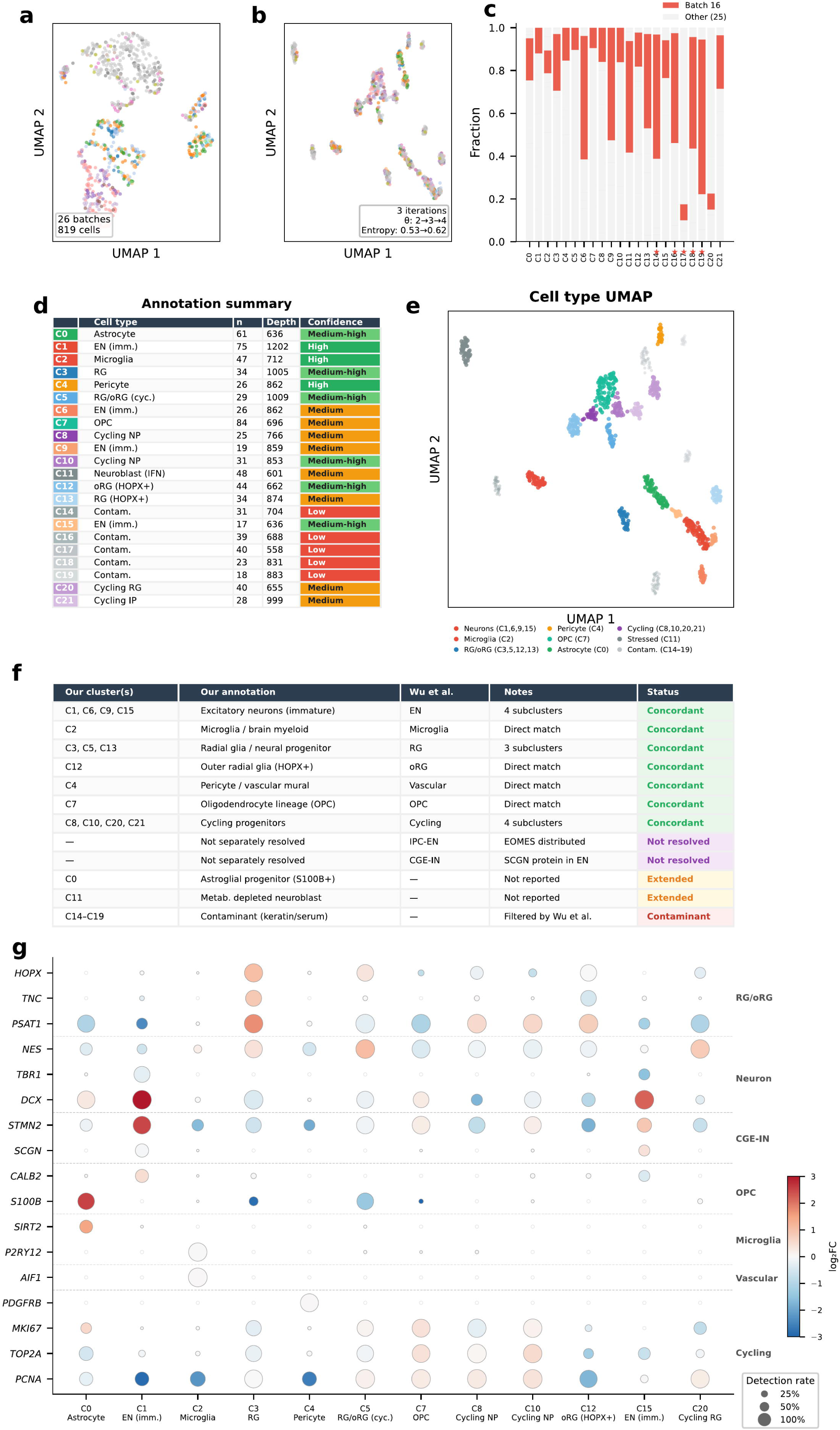
Benchmarking against the developing human brain single-cell proteome (Wu et al., Nat Biotechnol 2026). (**a**) UMAP projection of 819 cells coloured by batch (26 batches) before Harmony batch correction. Strong batch-driven structure is evident. (**b**) UMAP after adaptive Harmony correction (3 iterations; θ: 2.0→3.0→4.0; Shannon entropy: 0.53→0.62). Batches are well mixed and biological clusters emerge. (**c**) Cluster-per-batch composition. Red bars indicate the fraction of cells from batch 16 (the largest batch, n = 210); grey bars show all other batches combined. Stars mark clusters annotated as contaminants (C14-C19), which are predominantly batch-16-derived. (**d**) Annotation summary table for all 22 clusters, showing cell type, number of cells, median proteome depth, and confidence tier. (**e**) UMAP coloured by cell type annotation, with cluster colours matching the summary table. Contaminant clusters (grey) are plotted at reduced opacity. Legend groups clusters by broad lineage. (**f**) Concordance table mapping our 22 clusters to the 8 major cell types reported by Wu et al. Fifteen clusters are concordant across 6 of 8 major lineages; IPC-EN and CGE-IN are not separately resolved (see text); 2 clusters extend the published annotation (C0: astroglial progenitor; C11: metabolically depleted neuroblast); 5 clusters are contaminants. (**g**) Marker dotplot for 18 canonical brain cell type markers across 12 representative clusters. Dot size encodes detection rate; dot colour encodes intensity log₂ fold-change (red = enriched, blue = depleted).

Following correction, Leiden clustering identified 22 clusters (Fig. 2d & e), 15 of which mapped concordantly to 6 of the 8 major cell types described by Wu et al. (Fig. 2f). Excitatory neurons, microglia, radial glia, outer radial glia, vascular mural cells, oligodendrocyte-lineage cells and cycling progenitors were all recovered with the expected canonical markers (Fig. 2g). Two published cell types were not resolved separately. Intermediate progenitor cells marked by EOMES were distributed across cycling populations, and CGE interneuron markers such as SCGN and CALB2 were detected within excitatory neuron clusters rather than defining a distinct population. This is consistent with the original study’s observation that SCGN protein is less cell-type-specific than its transcript, and illustrates a genuine resolution limit of SCP for populations defined by transcript-protein discordant markers.

The brain dataset also exposed two annotation errors that proved informative. One cluster enriched for S100B and SIRT2 but lacking HOPX or TNC was labelled by the LLM as “astrocyte”, even though mature astrocytes are not expected at gestational weeks 13-19; “astroglial-progenitor” is the more appropriate designation. Another cluster showing broad depletion of biosynthetic machinery was labelled as an interferon-stressed neuroblast on the basis of weak DDX58 detection without broader programme support. These errors pointed to two recurrent failure modes: context-insensitive use of marker vocabulary and over-interpretation of sparse mechanistic signals.

We next applied the pipeline to the glioblastoma tumour-associated neutrophil (TAN) dataset of Sadiku & Brenes et al. ^12^, which presents a more difficult challenge because all cells belong to a single lineage. Here, the task is to distinguish functional states within neutrophils rather than separate major lineages, in a setting where neutrophil granule proteins are detected across almost all cells. In this context, detection rate alone becomes largely non-informative for many of the most abundant proteins, and interpretation depends heavily on integrating intensity-based and model-based evidence.

The adaptive Harmony loop converged after two iterations, increasing mean entropy from 0.55 to 0.67 (Fig. 3a-c). Leiden clustering identified nine clusters (3 d, e), four of which mapped to published functional states: activated TANs corresponding to a subset of Sadiku et al.’s Armed state, FCN1-positive inflammatory neutrophils at the Armed/Engaged boundary, OLFM4-positive degranulating cells matching the Vital Neutrophil Extracellular Traps (NETs) state, and a less-activated population corresponding to Exhausted neutrophils (Fig. 3e, f). The most revealing result, however, came from the discordant clusters. A protein-depleted cluster enriched for complement and coagulation factors was labelled as ambient-dominated debris but corresponds to the lytic NETosis population. A second cluster dominated by keratins and desmosomal proteins was called an epithelial contaminant but matches the immunosuppressive and angiogenic state and likely reflects phagocytic uptake of epithelial material. A third cluster combined mitochondrial and ribosomal enrichment with tissue-derived signal and partially matched the vascular immature state. Our clustering at resolution 0.8 fragments Sadiku et al.’s states differently than their resolution 1.45. Sadiku’s Armed state (74 cells, 27%) is larger than any single cluster in our analysis; C0 (47 cells) captures a subset, with additional Armed cells likely distributed across C2 and C3. The proportionality of other mappings is more consistent: C6 (34 cells) maps well to Vital NETs (33 cells, 12%), C1 (35 cells) to Exhausted (47 cells, 17%), C4 (16 cells) to Immunosuppressive & Angiogenic (23 cells, 8%), and C8 (21 cells) to Vascular Immature (22 cells, 8%).

**Figure 3.**
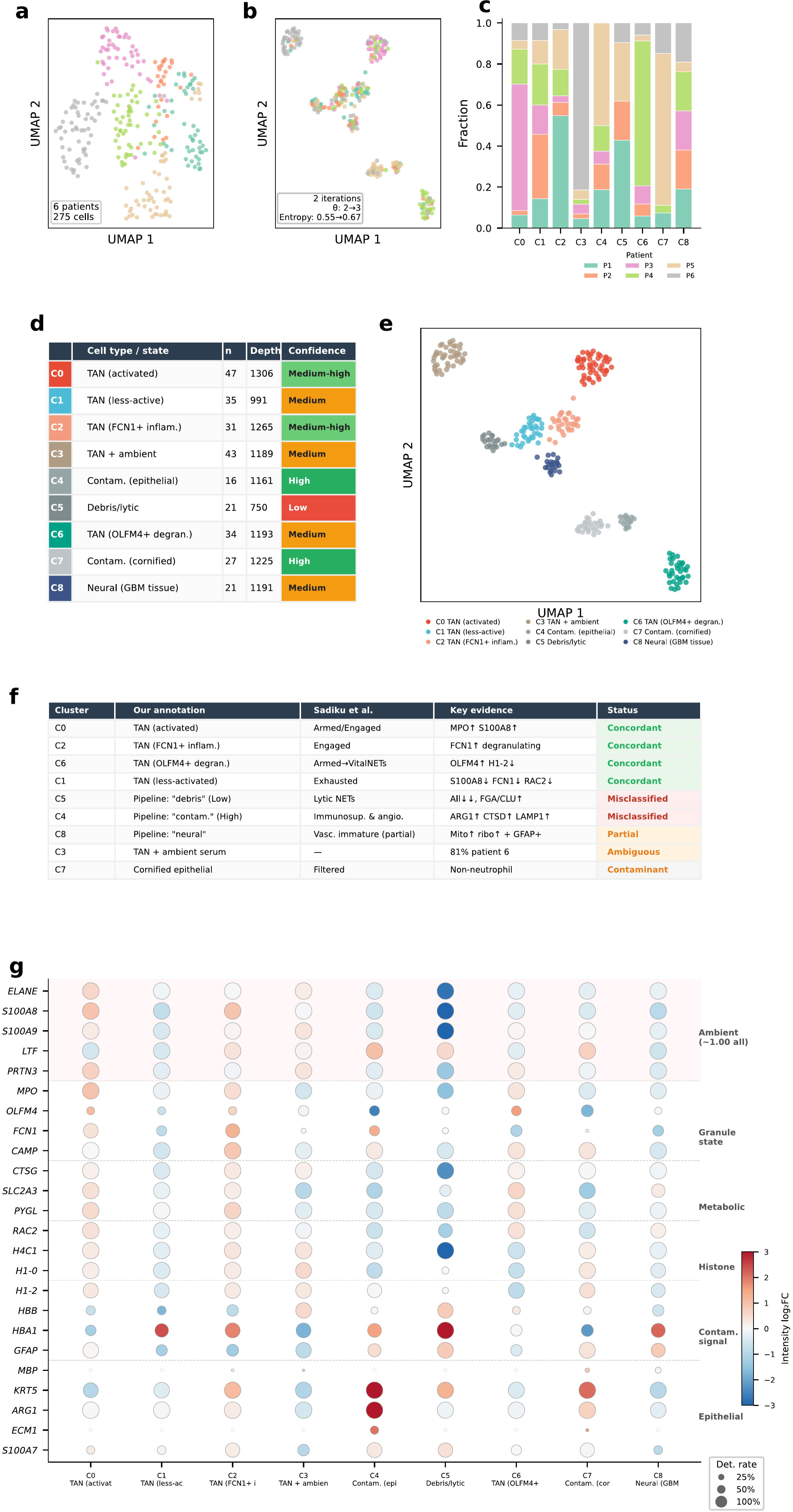
Benchmarking on glioblastoma tumour-associated neutrophils (Sadiku et al., Nat Commun 2026). (**a**) UMAP of 275 cells coloured by patient (6 GBM patients) before Harmony batch correction. Patient-driven structure is evident. (**b**) UMAP after adaptive Harmony correction (2 iterations; θ: 2.0→3.0; Shannon entropy: 0.55→0.67). (**c**) Cluster-per-patient composition. Each colour represents one patient. C3 is 81% patient 6; C6 is 71% patient 4; C7 is 74% patient 5. (**d**) Annotation summary table showing functional state, cell count, median proteome depth, and confidence tier. (**e**) UMAP coloured by annotated functional state. (**f**) Concordance table mapping our 9 clusters to Sadiku et al.’s 7 functional states. Four clusters are concordant; 2 are misclassified (C5 lytic NETs called “debris”; C4 immunosuppressive & angiogenic called “contaminant”); C8 partially matches vascular immature; C3 is ambiguous (patient-dominated); C7 is a genuine contaminant. (**g**) Marker dotplot for 27 proteins across all 9 clusters. The top 6 rows (red shading) demonstrate the ambient granule protein challenge: ELANE, S100A8/A9, LTF, PRTN3, and MPO are detected at ∼1.00 in ALL clusters including contaminants. Dot size (detection rate) is non-discriminative; only dot colour (intensity log₂FC) distinguishes activated from depleted neutrophils. Lower rows show markers where both detection rate and intensity provide discriminating information.

These failures shared a common interpretive logic: the model treated non-self-proteins primarily as evidence of contamination, rather than considering biologically plausible alternatives such as phagocytic uptake, vascular contact or lytic cell death. The neutrophil dataset, therefore, revealed that generic contamination rules are insufficient in cell states where uptake of extracellular material or proteome depletion is itself a defining biological feature.

### A three-round context-aware annotation architecture improves interpretation of ambiguous proteomic states

To address the failure modes uncovered by benchmarking, we revised the annotation framework. Structured prompting with explicit contradiction reasoning, confidence criteria and resolving-marker nomination improved several calls relative to an earlier unstructured prompt, but did not fully resolve the developmental and phagocytic-state errors identified in the brain and neutrophil datasets. We therefore introduced three additional principles: stage-appropriate vocabulary, a minimum evidence threshold for mechanistic labels, and explicit consideration of non-self protein acquisition mechanisms before assigning contamination.

To make these constraints generalisable rather than dataset-specific, we implemented a three-round annotation architecture (Fig. 4a,b). In Round 0, the model receives only the experiment context and is asked to infer likely cell types, vocabulary restrictions, anticipated ambient signals and plausible mechanisms of non-self-protein acquisition, without exposure to any cluster-level marker data. In Round 1, cluster annotation is performed using these inferred constraints together with the full marker summaries. In Round 2, clusters with low or medium confidence are revisited after extraction of supplemental markers nominated by the model itself. This design aims to constrain interpretation from the experimental setting rather than from reverse engineering of the observed results.

**Figure 4.**
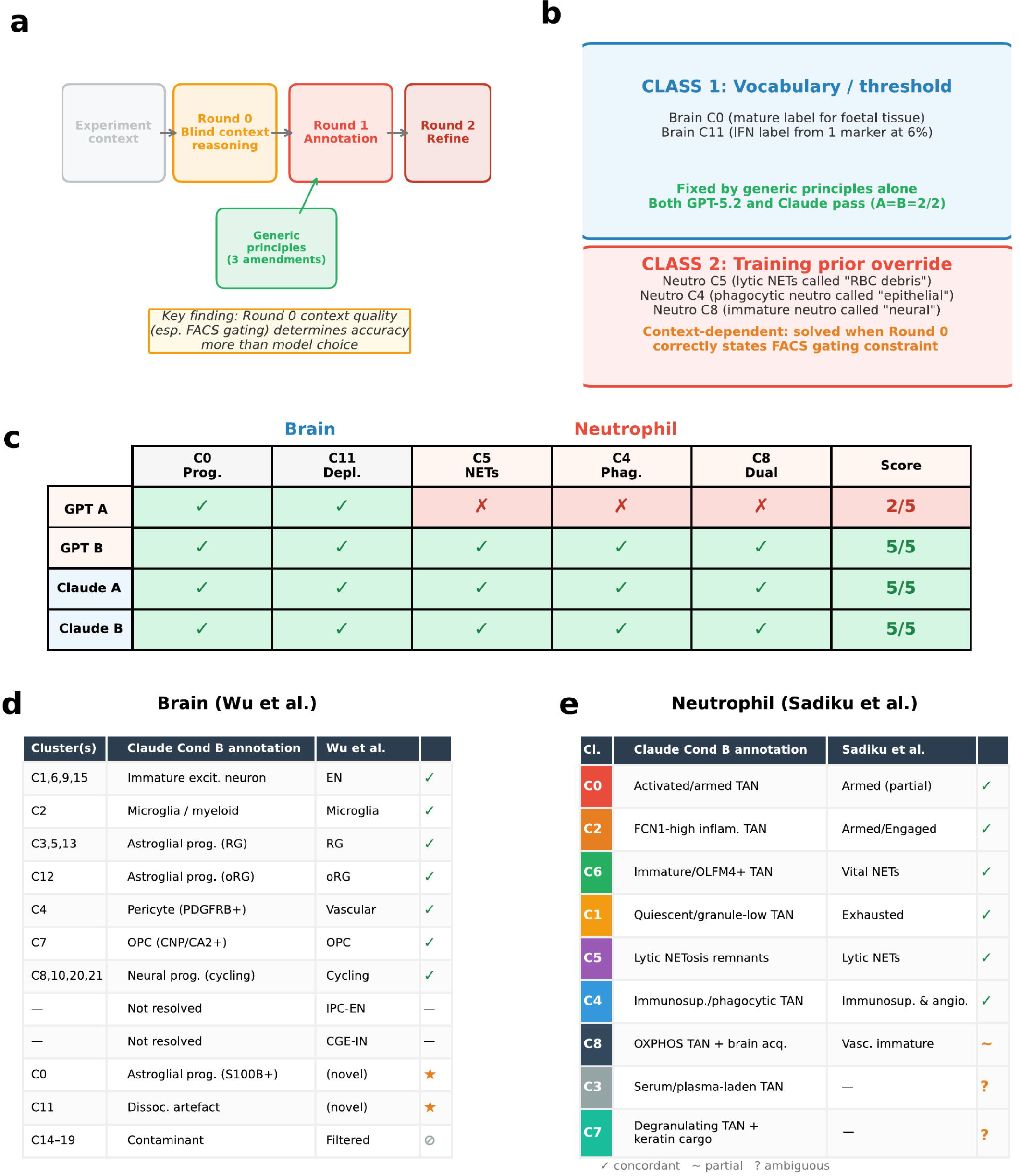
Prompt engineering architecture and model comparison. (**a**) Two classes of LLM annotation failure. Class 1 (vocabulary/threshold): developmental vocabulary and mechanistic annotation threshold errors, correctable by generic prompt principles. Class 2 (training prior override): failures requiring the LLM to override strong pattern-matching associations using experiment-context reasoning. (**b**) Round 0: the LLM generates dataset-specific analytical constraints from the experiment context alone, without seeing any cluster data. Round 1: clusters are annotated using generic amendments plus Round 0 constraints plus full marker data. Round 2: supplemental markers are queried for Low/Medium-confidence clusters. Three conditions were tested: A (generic principles only), B (generic + Round 0) (**c**) Heatmap showing annotation accuracy across 5 failure-mode targets (2 brain, 3 neutrophil) for GPT-5.2 and Claude Sonnet 4.6 under all three conditions. Brain targets are solved by generic principles in both models. Neutrophil targets show a model-dependent split: With FACS-corrected context, both models achieve 5/5 under Condition B. Without FACS context, GPT-5.2 achieves only 2/5, requiring hardcoded rules (Condition C) to reach 5/5; Claude achieves 5/5 regardless. (**d**) Brain concordance table: Claude Condition B annotations for all 22 clusters mapped against Wu et al.’s 8 cell types. Six of 8 types concordant; IPC-EN and CGE-IN not separately resolved; 2 novel populations identified; 5 contaminant clusters. (**e**) Neutrophil concordance table: Claude Condition B annotations for all 9 clusters mapped against Sadiku et al.’s 7 functional states. Seven of 7 states concordant with FACS-corrected context (C4 phagocytic and C5 lytic NETosis now correctly identified); C8 partially matches Vascular Immature; C7 reclassified from contaminant to degranulating TAN with keratin cargo.

Across the datasets examined here, Round 0 captured the key contextual constraints needed for improved interpretation. It correctly inferred, for example, that mature astrocytes are not expected in foetal cortex, that neutrophil granule proteins are likely to appear as ambient signal, and that phagocytosis or lytic cell death can plausibly explain non-self or protein-depleted proteomes. Under this framework, the developmental vocabulary and mechanistic-threshold errors observed in the brain dataset were resolved.

Model comparison nevertheless revealed a clear distinction between failures that are solvable by principle-based prompting and those that require stronger prior override. Both GPT-5.2 and Claude Sonnet 4.6 LLM corrected the brain targets under the general prompting conditions, indicating that these are largely problems of vocabulary control and evidential discipline. The neutrophil targets showed some model dependence (Fig. 4c-e). When the experiment context correctly stated that all cells are FACS-sorted neutrophils, both Claude and GPT-5.2 resolved all five failure-mode targets under Condition B. Without explicit FACS gating information, GPT-5.2 and Claude gave ambiguous results, demonstrating that the quality of the Round 0 experiment context is an important constraint. The difference was most evident in phagocytic clusters dominated by keratin cargo: without the FACS gating constraint, GPT-5.2 reverted to contaminant labels when faced with strong epithelial signatures despite correctly describing phagocytosis as plausible in Round 0. With the corrected context explicitly stating that all cells passed a CD45+CD66b+CD49d− neutrophil gate, both models correctly identified C4 as phagocytic and C5 as lytic NETosis.

### Held-out validation demonstrates cross-platform generalisability and agreement with orthogonal ground truth

We next tested whether the pipeline generalises to data not used during development by analysing a CYLD cutaneous syndrome (CCS) skin tumour ^17^ dataset acquired on a different instrument and using a different sample preparation workflow ^18^. This dataset provides a particularly strong validation setting because each cell has an orthogonal FACS sort label of two distinct cell types, allowing direct assessment of annotation accuracy at single-cell level.

Using default parameters and the three-round architecture, adaptive Harmony converged after a single iteration and Leiden clustering identified seven clusters (Fig. 5a,b). Claude annotated two clusters as keratinocyte subtypes and five as myeloid subtypes, including inflammatory macrophages, tissue-resident macrophages, stressed antigen-presenting myeloid cells and a distinct phagocytic macrophage population. Overall, 226 of 249 cells, corresponding to 90.8%, were assigned to clusters whose label matched the FACS sort identity, and six of the seven clusters showed at least 91% FACS purity (Fig. 5c,d).

**Figure 5.**
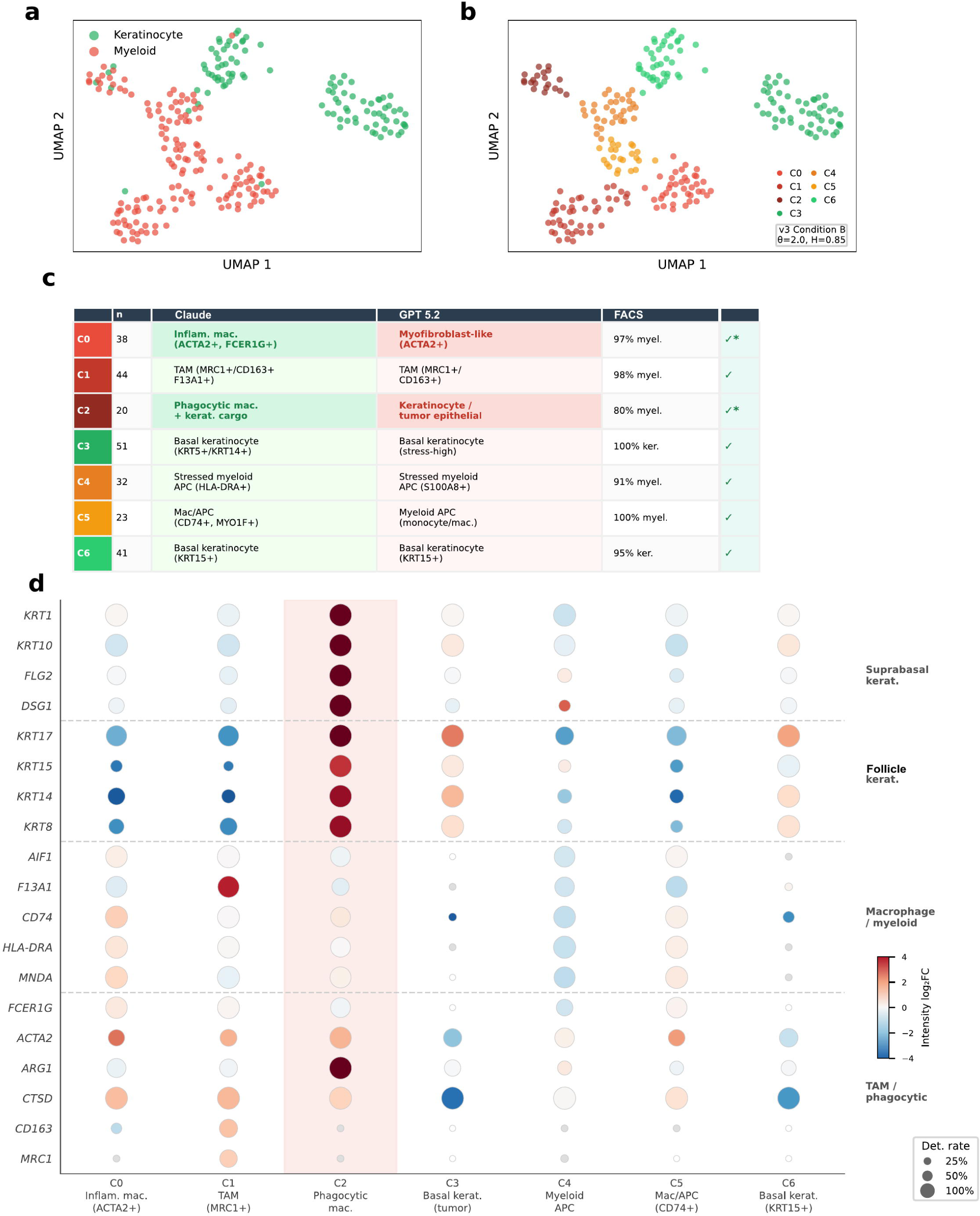
Held-out validation on CCS skin tumour with FACS ground truth (Inns et al. 2026). (**a**) UMAP of 249 cells from CYLD cutaneous syndrome (CCS) skin tumour coloured by FACS sort identity: CD45⁻/CD200⁺ keratinocytes (green) and CD45⁺/CD74⁺ immune cells (red). Data acquired on Bruker timsTOF HT with oil-immersion sample preparation - a different instrument and workflow from all development datasets. (**b**) UMAP coloured by pipeline clusters (7 clusters, Leiden resolution 0.8). Harmony converged in 1 iteration (θ=2.0, entropy=0.85). (**c**) Concordance table mapping v3/Condition B annotations (Claude) to FACS ground truth and Inns et al.’s SingleR cell type categories. FACS purity is ≥91% for 6 of 7 clusters; overall cell-level concordance: 226/249 = 90.8%. C2 comprises 80% CD45⁺/CD74⁺-sorted cells with dominant keratinocyte cargo (KRT1/KRT10/FLG2 log₂FC=+7.9); Claude v3 correctly identifies these as phagocytic macrophages while GPT-5.2 v3 calls them keratinocytes - replicating the model-dependent prior-override pattern from the neutrophil dataset. C1 (tissue-resident macrophage, CD163 det=0.68, MRC1 det=0.64) replicates Inns et al.’s key finding of CD163⁺/MRC1⁺/CD209⁺ TAMs. (**d**) Marker dotplot for 21 keratinocyte and myeloid markers across all 7 clusters. Suprabasal markers (KRT1, KRT10, FLG2, DSG1) are massively enriched in C2 (blue highlight); basal markers (KRT5, KRT14, KRT15) define C3 and C6. Myeloid markers (AIF1, F13A1, CD74, HLA-DRA, MNDA, CTSC) are enriched in C0/C1/C4/C5 and absent from C3/C6. TAM markers CD163 and MRC1 are specifically detected in C1. ARG1 is enriched in C2, consistent with keratinocyte-derived arginase rather than myeloid ARG1.

The most informative population was cluster C2. Although suprabasal and follicular keratinocyte proteins dominated its proteome, 80% of its cells originated from the CD45-positive, CD74-positive immune sort gate, and myeloid markers remained clearly detectable. Claude interpreted this pattern as macrophages carrying ingested keratinocyte cargo, consistent with tumour-associated phagocytosis. GPT-5.2, given the same context and marker evidence, instead labelled the cluster as keratinocyte or tumour epithelial, demonstrating discordance between the models.

A second informative case was cluster C0, which showed ACTA2 and VIM enrichment compatible with either myofibroblastic stromal cells or contractile macrophages. Because the experiment context specified that all cells had been sorted into either immune or keratinocyte gates, fibroblasts should have been excluded by design. Claude derived this constraint from the experimental description and annotated the cluster as inflammatory macrophages, whereas GPT-5.2 continued to return a myofibroblast-like label. These examples indicate that successful annotation in ambiguous proteomic states depends not only on marker interpretation, but also on the ability to infer exclusion constraints from experimental design.

Relative to the published SingleR-based analysis, which identified five broad cell types ^18^, our framework possibly resolved finer heterogeneity, including two keratinocyte subtypes, four myeloid subtypes and a phagocytic macrophage state that reference-based approaches would probably misclassify according to the dominant cargo-derived proteomic signature. Taken together, this held-out validation indicates that the pipeline generalises across instruments, workflows and tissues while maintaining agreement with independent experimental labels.

### Orthogonal tissue validation supports annotations in caerulein-injured pancreas

Finally, we applied the pipeline to mouse pancreas collected two days after caerulein-induced acute pancreatitis, a setting chosen because it combines epithelial injury, stromal activation and immune infiltration within a single dataset ^19^. After adaptive quality control, 234 cells were retained from four batches. Batch effects were modest, and the Harmony loop converged after a single iteration with high entropy, yielding eight clusters without evidence of batch-dominated structure (Fig. 6a,b).

**Figure 6.**
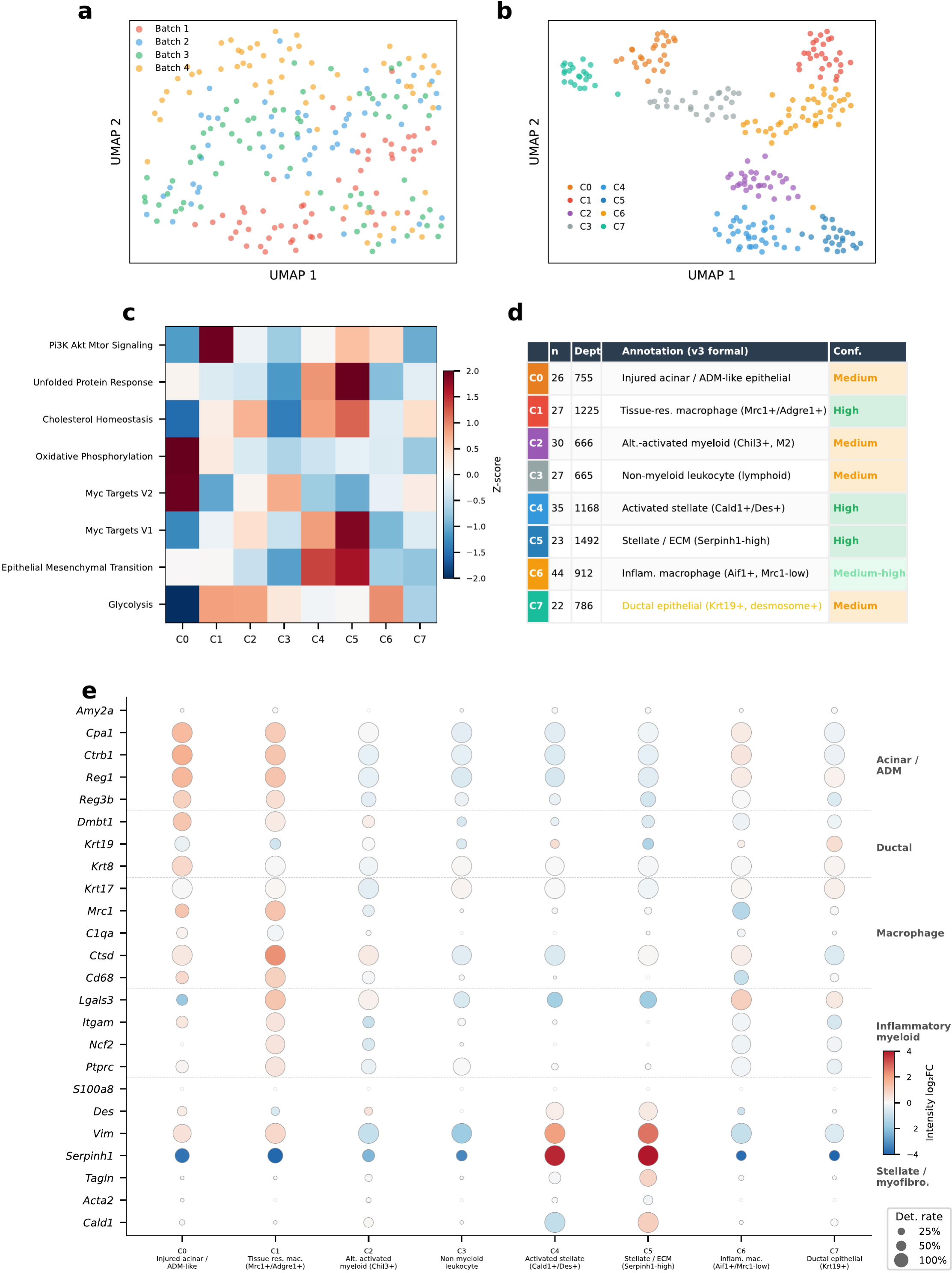
Pipeline validation with v3 annotation architecture on caerulein-injured mouse pancreas. (**a**) UMAP projection of 234 cells coloured by batch (4 libraries) before Harmony batch correction. No batch-driven or biological cluster structure is evident in the uncorrected embedding. (**b**) UMAP after adaptive Harmony correction (θ=2.0, Shannon batch mixing entropy=0.90, single iteration). 234 cells resolve into 8 clusters. (**c**) AUCell pathway activity heatmap showing the 8 most variable MSigDB Hallmark pathways across clusters (C0-C7), Z-scored per pathway. PI3K/AKT/mTOR signalling and unfolded protein response mark C0 (ADM-like epithelial, consistent with ER stress in injury-driven acinar remodelling); cholesterol homeostasis marks C1 (macrophage); epithelial-mesenchymal transition marks C4/C5 (stellate/myofibroblast); glycolysis marks C6 (inflammatory myeloid). (**d**) Annotation summary table showing v3/Condition B labels. Cluster colour badges (left column) match panel (b). Confidence tiers are colour-coded: green (High), light green (Medium-high), orange (Medium), red (Low). Depth = median detected proteins per cell. Key v3 improvements: C0 annotated as “ADM-like epithelial” rather than “Acinar” (Round 0 context constrains vocabulary to injury terminology); C1 macrophage confirmed at High confidence with Round 0 correctly noting acinar enzymes as phagocytic cargo; C4 upgraded to High confidence with Round 0 confirming activated stellate cells expected post-caerulein; C6 refined to “neutrophil-leaning inflammatory” with Round 0 noting neutrophils as first responders. (**e**) Marker dotplot for 24 key proteins grouped by 5 lineage categories (Acinar/ADM, Ductal, Macrophage, Inflammatory myeloid, Stellate/myofibroblast). Dot size encodes detection rate; dot colour encodes intensity log₂ fold-change (red = enriched, blue = depleted). Acinar enzymes (Amy2a, Cpa1, Ctrb1) are detected ubiquitously due to ambient carryover but show intensity enrichment only in C0, demonstrating why detection specificity alone is insufficient. Reg1/Reg3b (injury-regeneration markers) are enriched in C0, supporting the ADM-like designation. Macrophage markers (Mrc1, C1qa, Ctsd) are specific to C1. The stellate activation gradient is visible: Des marks both C4 and C5, while Serpinh1 shows the strongest enrichment in C5 (myofibroblast, median 1,492 proteins -highest proteome depth in the dataset).

The resulting clusters spanned the major cellular compartments expected in the injured pancreas. Pathway-level analysis revealed distinct functional programmes across clusters (Fig. 6c). C0 showed the strongest oxidative phosphorylation enrichment, consistent with high metabolic demand of injured cells. C1, C2,and C6 were enriched for Glycolysis whereas EMT and the unfolded-protein response were enriched across both C4 and C5 (Fig. 6c). Taking molecular and pathway markers, the annotation framework assigned high confidence to tissue-resident macrophages and two stellate-lineage populations, medium-high confidence to an inflammatory macrophage population and medium confidence to four additional clusters representing injured epithelial, alternatively activated myeloid, low-depth lymphoid-leaning and ductal-like populations (Fig. 6d,e).The lymphoid-leaning cluster is likely to be a mis-annotation as it resembles an acinar-like population with low proteome coverage, likely to be dead or dying cells of acinar origin.

This dataset also reinforced the value of multi-modal interpretation. Acinar enzymes were detected broadly across clusters because of ambient digestive-enzyme carryover, but showed intensity enrichment only in the injury-remodelled epithelial population, making detection alone insufficient for annotation. Likewise, the stromal compartment resolved into a stellate activation continuum rather than a single fibroblast-like class, with Des marking lineage identity and Serpinh1, Tagln and Acta2 marking increasing activation. Importantly, the framework also withheld stronger claims where evidence was incomplete, nominating additional lineage markers for the shallow lymphoid-leaning cluster and maintaining low confidence for the ductal-like cluster in the absence of canonical ductal markers, suggesting that both these clusters require additional investigation.

To provide non-computational support for these annotations, we performed immunohistochemistry and immunofluorescence on FFPE sections from the same caerulein-injured pancreas tissue (Fig. 7). Six markers were selected to span the four major compartments identified by the pipeline. CD68 confirmed macrophage infiltration within the injured parenchyma, with positive cells distributed between acinar structures in a pattern consistent with the C1 tissue-resident macrophage population (Fig. 7a). Galectin-3 (Lgals3), detected at 100% in C1 and C6 by SCP with significant intensity enrichment (log_2_FC = +1.23 and +0.98, respectively), showed strong staining in interstitial cells morphologically consistent with macrophages, with weaker but detectable signal in acinar cells, matching the lower detection rate observed in C0 (31%) (Fig. 7a). Amylase staining appeared patchy across the injured tissue, consistent with heterogeneous loss of digestive enzyme expression during acinar-to-ductal metaplasia. In the SCP data, Amy2a5 was detected ubiquitously due to ambient carryover but showed intensity enrichment restricted to C0 (log₂FC = +0.93), underscoring the complementarity of tissue-level and single-cell proteomic readouts (Fig. 7a). Krt17/19 staining was localised to ductal structures, supporting the C7 ductal-like annotation (Fig. 7a). The stellate activation gradient inferred from SCP was corroborated by differential staining of Vimentin and Tagln: Vimentin was broadly expressed in stromal cells and some interstitial immune cells, consistent with its significant intensity enrichment in both C4 (log_2_FC = +2.06) and C5 (log_2_FC = +2.71), whereas Tagln staining was more restricted, in keeping with its enrichment predominantly in C5 myofibroblasts (70% detection) over C4 stellate cells (34% detection) (Fig. 7a).

**Figure 7.**
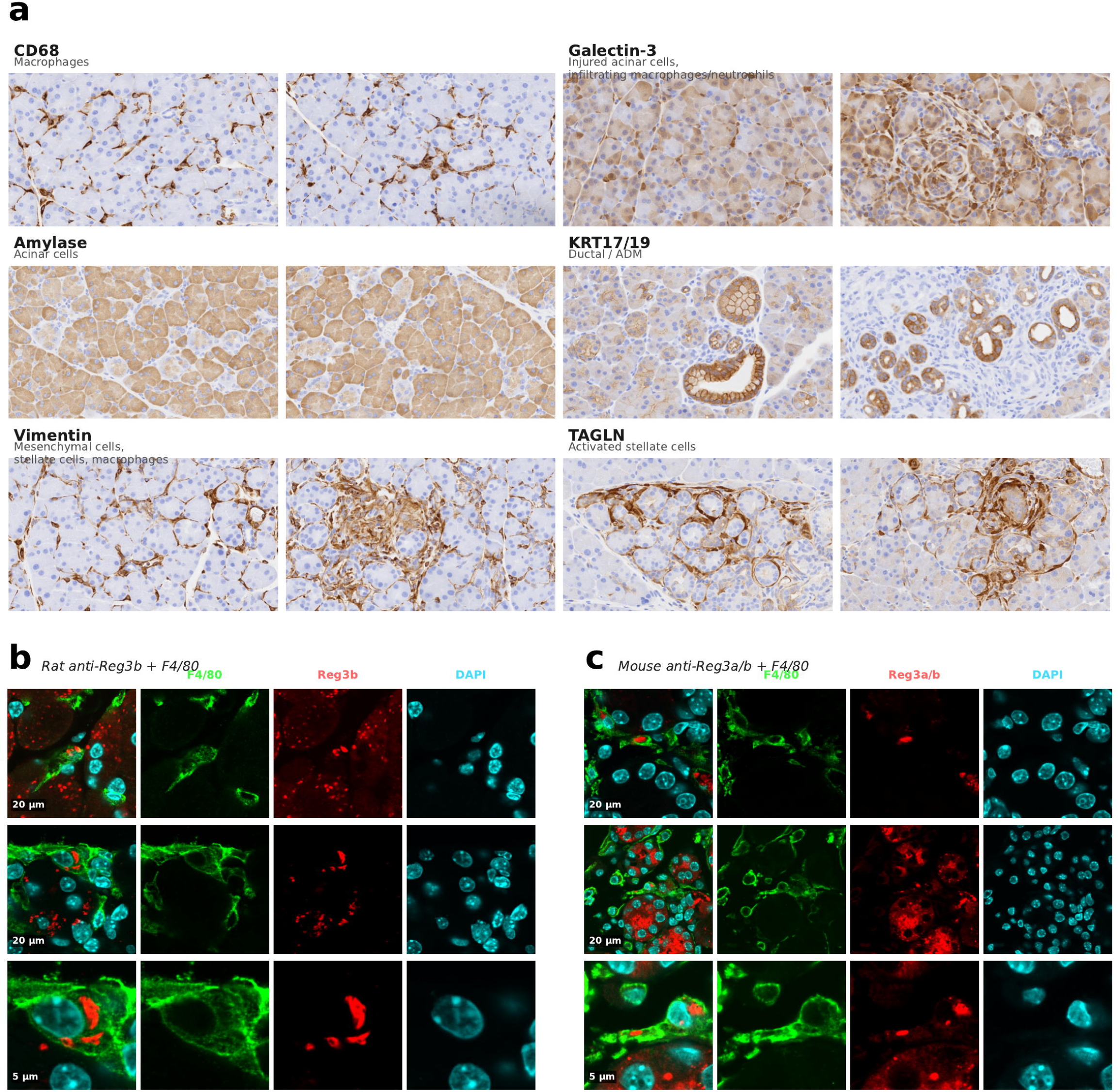
Orthogonal tissue validation of pipeline annotations in caerulein-injured mouse pancreas. (**a**) Immunohistochemistry on FFPE sections from the same injured pancreas tissue used for SCP. Six markers validate the four major compartments identified by the pipeline. Two representative fields per marker. (**b**) Immunofluorescence co-staining for Reg3b (anti-Reg3b antibody, red) and F4/80 (macrophage marker, green) with DAPI nuclear counterstain (cyan). Top row: two 20 µm overview fields showing Reg3b signal within and adjacent to F4/80-positive macrophages in the injured parenchyma. (**c**) Immunofluorescence co-staining for Reg3a/b (anti-Reg3a/b antibody, red) and F4/80 (green) with DAPI (cyan), confirming the co-localisation pattern with an independent antibody. Layout as in (b). Scale bars: 20 µm (overview) and 5 µm (zoom).

To directly test the pipeline’s inference that macrophage-associated Reg3b signal reflects phagocytic uptake rather than ambient contamination, we performed immunofluorescence co-staining for Reg3 protein and F4/80 (a macrophage surface marker) on adjacent sections (Fig. 7b,c). Reg3b was detected in 82% of C1 macrophages by SCP, with modest intensity enrichment (log_2_FC = +0.60). In tissue sections, Reg3 protein was visible as discrete intracellular puncta within F4/80-positive macrophages, consistent with phagocytic uptake of acinar-derived Reg3 protein. This pattern was confirmed using two independent antibodies, a rat anti-Reg3b and a mouse anti-Reg3a/b, both showing co-localisation with F4/80 at 20 µm overview and 5 µm high-magnification scales. These results provide direct visual evidence that non-self acinar proteins detected in the macrophage proteome by SCP originate from ingested cellular material, supporting the pipeline’s interpretation and reinforcing the broader principle that non-self protein acquisition should be considered before assigning contamination labels in phagocytic cell populations.

## Discussion

We present a fully automated end-to-end pipeline for single-cell proteomics analysis that integrates adaptive quality control, iterative batch correction, multi-modal marker discovery and context-aware annotation. Across four datasets spanning different tissues, instruments, species and analytical challenges, the framework achieved substantial concordance with published expert annotations and independent experimental labels, while making its own uncertainty and failure modes explicit.

A central conclusion from these analyses is that multi-modal evidence integration is essential for SCP. In relatively straightforward lineage-resolution tasks such as the foetal brain dataset, detection patterns and canonical markers are often sufficient to recover major populations. In more difficult settings, such as tumour-associated neutrophils or injured pancreas, however, pervasive ambient signal makes detection rate alone unreliable. Here, intensity fold-change, model-based effects and pathway-level signatures become critical for distinguishing genuine biological enrichment from broadly distributed background. The held-out skin tumour dataset, analysed across a different instrument and sample preparation workflow, suggests that this multi-modal strategy generalises beyond the development datasets used to build the pipeline.

The LLM annotation layer adds a different kind of capability: the ability to weigh contradictory evidence, incorporate experimental context and explicitly communicate uncertainty. At the same time, our evaluation identified two broad classes of failure. The first comprises vocabulary and threshold failures, such as assigning adult cell-type language to developmental populations or inferring mechanistic states from sparse evidence. These errors were largely corrected by general prompt principles. The second comprises prior-override failures, in which the model must reject a strong pattern-matching interpretation in favour of a contextually grounded explanation. Examples include phagocytic neutrophils or macrophages whose proteomes are dominated by acquired epithelial cargo, and lytic NETosis remnants whose depleted proteomes resemble debris. These cases proved substantially more model-dependent.

The three-round architecture provides a principled way to address such problems without embedding answers directly into the prompt. By deriving dataset-specific constraints from experiment context before exposing the model to marker evidence, the framework constrains interpretation using design-level knowledge rather than reverse-engineered outcomes. In practice, this approach transforms annotation uncertainty from a weakness into a productive output: contradictory markers are surfaced explicitly, low-confidence clusters trigger resolving- marker nomination, and alternative explanations can be carried forward into orthogonal validation.

Several limitations should be acknowledged. First, the annotation layer currently depends on commercial LLM APIs, introducing cost and availability constraints. Second, concordance with published studies was mainly assessed at cluster level because our clustering and resolution settings do not match those used in the original analyses. Third, the datasets examined here range from 234 to 819 cells; larger SCP datasets may reveal additional scaling constraints that are not captured in the present study. Fourth, Leiden resolution and dual-modality embedding were used as default choices and were not benchmarked exhaustively against alternative parameterisations or intensity-only embeddings. Finally, our comparison of model behaviour should be interpreted cautiously: the prompt architecture was developed iteratively in the course of this work, and broader blinded replication across additional model families will be needed before drawing strong conclusions about relative reasoning capacity.

More broadly, the phagocytic-cell problem highlighted here is likely to extend beyond LLM-assisted annotation. Any automated SCP framework that equates non-self-proteins with contamination risks systematic misclassification in biological contexts where uptake, lysis or extracellular exposure are central features of cell state. As SCP datasets continue to grow in scale and complexity, the combination of structured multi-modal evidence, context-aware reasoning, explicit uncertainty quantification and orthogonal validation offers a practical foundation for reproducible and interpretable analysis.

## Methods

### Pipeline implementation

The SCP analysis pipeline was implemented as a 22-rule Snakemake workflow (v7.32+) with full dependency resolution and checkpointing. All processing stages produce structured JSON metadata files for provenance tracking. The pipeline accepts protein group matrices from DIA-NN, Spectronaut, or FragPipe and produces quality-controlled, batch-corrected, clustered, and annotated outputs with per-cluster marker statistics across four complementary modalities.

### Datasets

Three published SCP datasets were used for validation and benchmarking. The caerulein-injured mouse pancreas dataset (D2_SCP) comprised 361 cells from 4 independently processed libraries, searched with Spectronaut against UniProt mouse proteome (UP000000589) using the mz_300_1200_frag_200_2000 preset, trypsin digestion (K*, R*), 2 missed cleavages, and no carbamidomethylation; 234 cells passed QC (≥500 detected proteins). The developing human brain dataset (Wu et al., Nat Biotechnol 2026) comprised 2,310 cells from prenatal cortex at gestational weeks 13, 15, and 19, searched with DIA-NN (direct FASTA mode) against UniProt human proteome (UP000005640) with acetyl N-terminal and oxidised methionine variable modifications and carbamidomethyl cysteine as fixed modification; 819 cells from 26 batches passed QC, with 4,475 protein groups identified. The glioblastoma tumour-associated neutrophil dataset (Sadiku et al., Nat Commun 2026) comprised 330 FACS-sorted CD45+CD66b+CD49d− neutrophils from 6 GBM patients, searched with Spectronaut; 275 cells from 6 batches passed QC, with 3,869 protein groups identified and a median of >1,100 proteins per cell. For held-out validation, the CYLD cutaneous syndrome skin tumour dataset (Inns et al., bioRxiv 2026) comprised 419 FACS-sorted cells from CCS tumours (CD45⁻/CD200⁺ keratinocytes and CD45⁺/CD74⁺ immune cells), acquired on a Bruker timsTOF HT with oil-immersion sample preparation and searched with DIA-NN v2.0.1 against UniProt human proteome with a neural-network-derived spectral library; 249 cells from 6 plates passed QC (≥400 detected proteins), with 5,934 protein groups identified and a mean of ∼700 proteins per cell. FACS sort labels were extracted from run metadata and used as orthogonal experimental ground truth.

### Quality control and cell filtering

The pipeline applies an adaptive cell filtering strategy. Library, carrier, and multi-cell control runs are excluded by configurable regex matching. Remaining cells are filtered using an adaptive cutoff: the median number of detected proteins across the bottom-N cells (default N=5), multiplied by a user-adjustable factor (default 1.7×), with a hard floor of 400 detected proteins. Six diagnostic plots are generated per dataset, including a cluster-per-batch composition metric that flags batches whose cluster distribution diverges from the global pattern.

### Dual-modality embedding and iterative batch correction

Joint cell embeddings are constructed from a dual-modality PCA that concatenates standardised principal components from (i) log₂-transformed, median-normalised protein intensities (20 PCs, median-imputed, z-scored) and (ii) binary detection patterns (10 PCs, centred). This design exploits the informative missingness unique to SCP: which proteins are detected is as informative as how much protein is measured. The pipeline implements an adaptive Harmony batch correction {Korsunsky, 2019 #16} loop: after each Harmony run, weighted Shannon entropy of batch mixing across Leiden clusters is computed (range 0-1, where 1 = perfect mixing). If mean entropy falls below the target threshold (default 0.6), the diversity penalty θ is incremented by 1.0 and correction is re-run. This continues until the entropy target is met or maximum θ is reached (default 5.0). The pancreas dataset converged in 1 iteration (θ=2.0, entropy=0.90); the brain dataset required 3 iterations (θ: 2.0→3.0→4.0, entropy: 0.53→0.57→0.62); the neutrophil dataset required 2 iterations (θ: 2.0→3.0, entropy: 0.55→0.67). Unsupervised clustering was performed using Leiden community detection at default resolution (0.8). UMAP was computed for visualisation using default parameters.

### Per-cluster marker statistics

Per-cluster differential marker statistics were computed across four complementary modalities. *Detection specificity*: Fisher’s exact test on binary protein presence/absence, comparing each cluster against all remaining cells, with Benjamini-Hochberg FDR correction (q<0.05). *Intensity fold-change*: Wilcoxon rank-sum test applied exclusively to cells in which the protein was detected (detected-only), avoiding zero-inflation confounding. *scplainer*: a linear mixed-effects model decomposing per-protein expression into biological (cluster) and technical (batch, total detected proteins, injection order) components. *Pathway activity*: AUCell applied to MSigDB Hallmark gene sets, computing per-cell AUC scores and mean scores per cluster. Proteins ranked concordantly across ≥2 modalities were designated consensus markers by combined Borda rank score (|fold-change| × −log_10_(q-value), min-max normalised per modality, averaged).

### Context-aware LLM annotation framework

Cell type annotation was performed using a structured prompt assembled from five components: (i) an experiment context block specifying species, tissue, experimental condition, and developmental stage; (ii) explicit definitions of each marker modality; (iii) a four-tier confidence rubric (High: ≥3 canonical markers concordant across ≥2 modalities, no contradictions; Medium-high: 2-3 markers, minor contradictions; Medium: 1-2 markers, contradictions present; Low: markers absent or contradicted); (iv) output format specifications requiring structured table with cell type, supporting markers, contradictions, confidence, and resolving markers; and (v) dataset-level caveats for proteins known to be ambient or non-discriminative.

Annotation proceeded in two sequential passes. In Pass 1, the complete per-cluster marker summary was submitted. Clusters assigned Low or Medium confidence triggered an automated supplemental marker query: proteins nominated by the model as resolving were extracted from the full protein group matrix and their per-cluster statistics appended. Pass 2 re-presented these clusters with supplemental evidence.

The prompt includes explicit contamination handling rules: the model is instructed not to label clusters as contaminants solely based on keratin, serum, or haemoglobin detection. Contamination classification requires that (a) no coherent lineage markers exist, (b) contaminant markers dominate >80% of top-ranked proteins, and (c) no alternative biological explanation is plausible.

### Three-round annotation architecture (v3)

Building on the two-pass framework, we developed a three-round architecture to generate dataset-specific constraints without hardcoding answers. In Round 0 (context reasoning), the LLM receives only the experiment context block - species, tissue, developmental stage, sample preparation - and generates analytical constraints covering: (i) vocabulary restrictions (cell types expected/absent at this developmental stage); (ii) anticipated ambient signals; (iii) non-self protein acquisition mechanisms relevant to the cell types in this experiment; and (iv) expected technical artefacts. Round 0 sees no cluster data, marker statistics, or annotation results - it reasons purely from the experiment description. Round 0 output is injected verbatim into the Round 1 annotation prompt alongside three generic prompt amendments: developmental context awareness, mechanistic annotation thresholds (≥2 concordant pathway markers at ≥10% detection), and non-self protein interpretation (phagocytosis, NETosis, trogocytosis, ambient carryover). Round 2 proceeds as the standard Pass 2 refinement.

### Model comparison

To assess model-dependence of annotation quality, the three-round architecture was tested under three conditions (A: generic amendments only; B: generic + Round 0; C: hardcoded rules with explicit thresholds) using two LLMs: GPT-5.2 (OpenAI) and Claude Sonnet 4.6 (Anthropic, claude-sonnet-4-6-20250219). All calls used temperature 0 for deterministic output. The exact model version, rendered prompt SHA-256 hash, and full raw output were recorded for each run. Five failure-mode targets were defined from the critical assessment in R3-R4: brain C0 (must use “progenitor” not “astrocyte”), brain C11 (must not use “IFN”; must mention depletion/artefact), neutrophil C5 (must identify NETosis), neutrophil C4 (must identify phagocytic neutrophil), and neutrophil C8 (must identify both neutrophil and tissue-derived signal). Regression was assessed on all three datasets by comparing v3 annotations with v2 annotations at the lineage level.

### Marker coverage score and PanglaoDB cross-validation

A marker coverage score was computed for each cluster as the proportion of recommended panel markers detected at ≥0.20 detection rate. Recommended panels were defined a priori for each expected cell type. Cross-validation against PanglaoDB (8,286 marker-cell type associations, 178 cell types) was performed by fuzzy-matching LLM-assigned labels to PanglaoDB cell types and computing the fraction of database-recommended markers detected.

### Reproducibility and audit trail

All LLM API calls used temperature 0. Prior to submission, rendered prompt text was hashed using SHA-256; hashes were stored alongside outputs in structured sidecar JSON files recording model version, timestamp, prompt character count, number of clusters, pass structure, and PanglaoDB usage. The pipeline is implemented as a Snakemake workflow with full dependency tracking; all intermediate outputs are versioned.

### Software and statistical analysis

The pipeline uses Python 3.10+ with scanpy (1.9+), harmony-pytorch, scikit-learn, scipy, and statsmodels. Visualisation uses matplotlib (3.7+). Marker statistics use scipy.stats (Fisher’s exact, Wilcoxon rank-sum) with statsmodels for multiple testing correction. AUCell pathway scoring uses the decoupler package with MSigDB Hallmark gene sets. All code is available at https://github.com/vonkriegsheim/CASPA.

### Animal work

All animal work was performed in accordance with institutional guidelines under license from the UK Home Office project license PP7280430 and approved by a University of Edinburgh internal ethics committee. Mouse was a 10-week-old female *Kras^LSL-G12D^* on a C57BL6/129Sv mixed background ^20^. 80ug/kg of caerulein (Bachem, 4030451) in PBS were administered by intraperitoneal (IP) injection once every hour for 6 hours. Dissociation was performed as described in Makar et al ^21^.

### Immunohistochemistry and immunofluorescence

Immunohistochemistry was performed on 4 µm FFPE sections from the same caerulein-injured mouse pancreas used for SCP. Sections were stained with antibodies against CD68 (CST), Galectin-3 (AF11, goat), Amylase (CST 3796, rabbit), Krt17/19 (CST 12434, rabbit), Vimentin (CST 5741, rabbit), and Tagln (Abcam ab14106, rabbit). Antigen retrieval was performed by immersing the slides in 10 mM sodium citrate, pH 6.0 and boiling them for 10 min using a microwave. Slides were then scanned using a Nanozoomer slide scanner (Hamamatsu Photonics) and viewed using NDP.view 2 software. For immunofluorescence, adjacent sections were co-stained with F4/80 (Cell Signaling Technology, D4C8V]) and either rat anti-Reg3b (R&D MAB5110-SP) or mouse anti-Reg3a/b (Santa Cruz, sc-377038). Nuclei were counterstained with DAPI. Images were acquired on an Olympus FV3000 Confocal Laser Scanning Microscope using Fluoview FV31S-SW (version 2.4.1.198) software and an Olympus UPlanSApo 60× 1.35 NA Oil objective.

### Pancreatic tissue sample processing for single-cell mass spectrometry

Single-cell processing was performed as previously described ^21^. Briefly, 1000 nL of master mix (0.2% DDM, 100 mM TEAB, 3 ng/μL trypsin) was dispensed into a 384-well plate (Eppendorf, 0030129547). The plate was then transferred into CellenONE for single-cell seeding. Cells were isolated based on isolation criteria: an elongation factor of up to 2.0 and a diameter range of 10−30 μm. After cell isolation, the plate was sealed with Thermowell sealing tape (Corning, 6569) followed by incubation for 2 h at 50 °C in a thermal cycler (PE Applied Biosystems, GeneAmp PCR System 9700). Post incubation the temperature was reduced to 20 °C, and 3.5 μL of 1% trifluoroacetic acid (TFA) was added to the samples.

Samples were then loaded onto purification and loading trap columns (Evosep, Evotips). Evotips were prepared according to manufacturer’s instructions. After loading the samples, the Evotips were washed with 0.1% FA followed by a final wash with 100 μL of 0.1% FA and spinning for 10 s at 800 g. The samples were then transferred into an Evosep One LC system for LC-MS/MS analyses.

### LC-MS/MS

LC–MS/MS analyses were performed using a timsTOF SCP mass spectrometer (Bruker) coupled to an Evosep One LC system (Evosep). Aurora Elite CSI analytical columns (IonOpticks; AUR3-15075C18-CSI) and a CaptiveSpray ionization source were used. Samples were analysed using the Whisper Zoom 40 SPD method, which employed a gradient flow of 100 nL/ min with a 31 min method duration. Eluted peptides were analysed with a parallel accumulation-serial fragmentation data-independent acquisition (diaPASEF) method. Method used for data acquisition consisted of 4 TIMS ramps with 5 mass ranges per ramp spanning from 327 to 1200 m/z and from 0.7 to 1.30 1/K0. Spectronaut 20 was used for data analysis, searching against the Uniprot Mus musculus database.

### Use of LLM statement

Claude code was used for coding de-bugging and linking python and R code using Snakemake, ChatGPT5.4 as embedded in the Edinburgh Language Model (ELM) was used to edit and correct grammar/flow of the manuscript draft.

## Data availability

The mass spectrometry proteomics data have been deposited to the ProteomeXchange Consortium with the dataset identifier PXD076177

## Acknowledgements

We are grateful to members of the von Kriegsheim laboratory for their discussions and critical reading of the manuscript. S.W. is supported by a CRUK Senior Fellowship (C20685/A29576); A.V.K. is supported by an MRC Grant (MR/X01293X/1) and a BBSRC Grant (BB/X019160/1). N.R and J.I. are supported by the NIHR Newcastle Biomedical Research Centre (BRC). AJB is supported by a Wellcome Trust Early Career Award (327847/Z/25/Z)

---

**Supplementary Figure 1. Detailed technical pipeline schematic.** (**a**) Complete directed acyclic graph (DAG) of the SCP analysis pipeline, showing all 22 Snakemake rules organised by execution level with data dependencies, parameters, and output files. (**b**) Key configurable parameters and their default values. (**c**) Statistical methods summary.

