## Supplementary material for "A Context-Aware Single-Cell Proteomics Analysis pipeline": Fig S1

### Supplementary Figure 1. Detailed technical pipeline schematic with DAG levels, parameters, and statistical methods

#### a Pipeline DAG — 22 rules across 10 execution levels

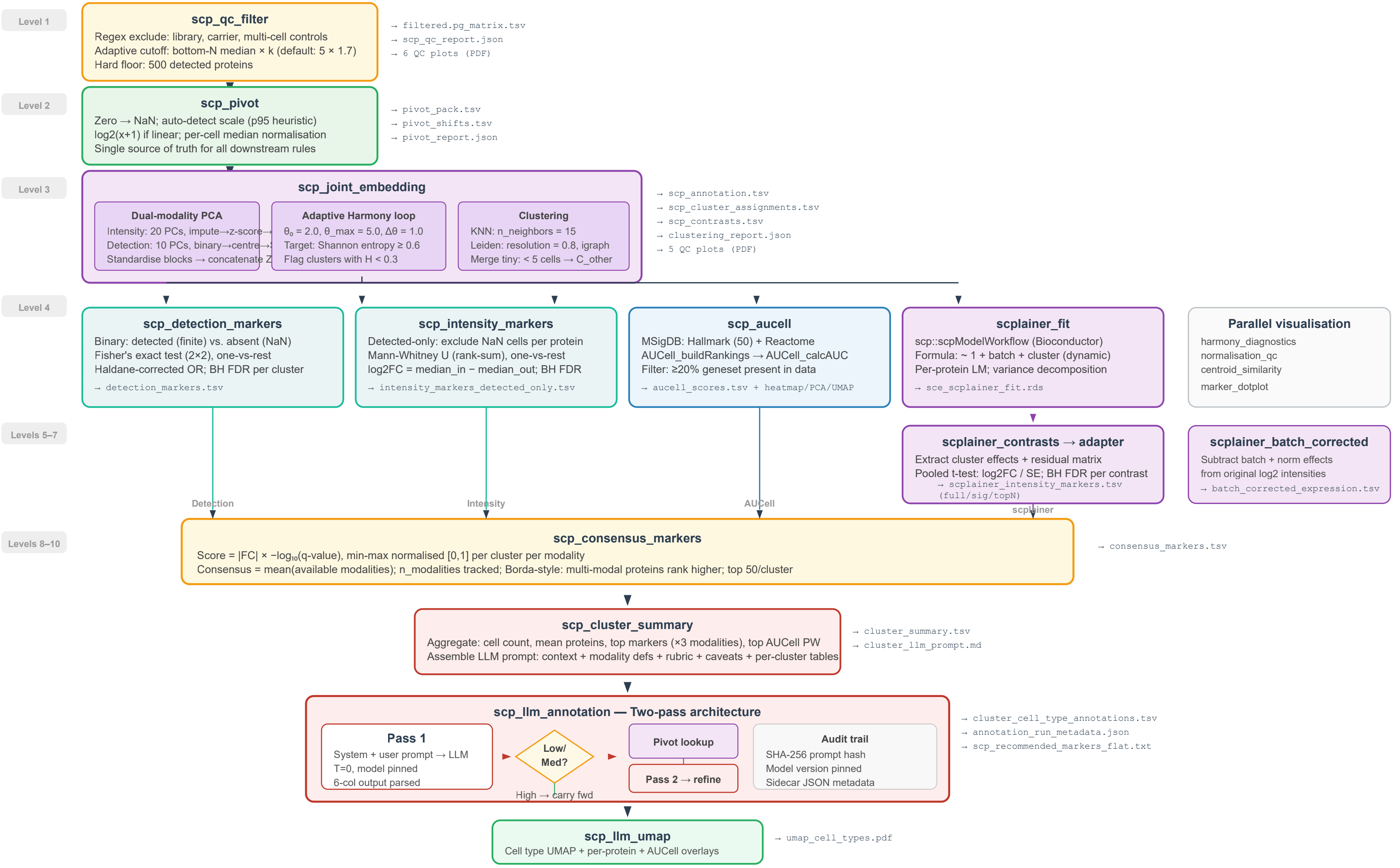


#### b Key configurable parameters and defaults

| Module | Parameter | Default | Description |
|---|---|---|---|
| QC | exclude_run_regex | ^(library\|lib\|5cell... | Regex for forced run exclusion |
| | min_protein_ids | 500 | Hard floor for detected proteins |
| | qc_bottom_n × qc_multiplier | 5 × 1.7 | Adaptive cutoff: median(bottom N) × k |
| Embed | n_pcs_int / n_pcs_det | 20 / 10 | PCs for intensity and detection blocks |
| | harmony_theta / theta_max | 2.0 / 5.0 | Diversity penalty range for adaptive loop |
| | harmony_entropy_target | 0.6 | Target batch mixing entropy (Shannon, weighted) |
| | n_neighbors / resolution | 15 / 0.8 | KNN graph + Leiden clustering resolution |
| Markers | min_cells_detected | 5 | Min cells for detection test (both modalities) |
| | consensus_top_n | 50 | Top markers per cluster in consensus |
| LLM | model / temperature | gpt-5.2 / 0 | Deterministic inference, model version pinned |

#### c Statistical methods

| Analysis | Test | Correction | Software |
|---|---|---|---|
| Detection markers | Fisher's exact (2×2) | BH FDR | scipy.stats.fisher_exact |
| Intensity markers | Mann-Whitney U | BH FDR | scipy.stats.mannwhitneyu |
| scplainer model | Per-protein LM | — | scp::scpModelWorkflow |
| scplainer DA | Pooled two-sample t | BH FDR per contrast | Custom R (matrixStats) |
| AUCell scoring | AUC on gene rankings | — | AUCell::AUCell_calcAUC |
| AUCell markers | Wilcoxon rank-sum | BH FDR | wilcox.test (R) |
| GO enrichment | Hypergeometric ORA | BH FDR | clusterProfiler::enrichGO |
| Consensus ranking | Borda count (\|FC\|×$-\log_{10}$q) | Min-max per modality | Custom Python |
| Batch correction | Harmony (iterative θ) | Entropy convergence | harmonypy.run_harmony |
| Clustering | Leiden community detection | — | scanpy.tl.leiden (igraph) |
